# Investigating the entropic nature of membrane-mediated interactions driving the aggregation of peripheral proteins

**DOI:** 10.1101/2022.01.24.477571

**Authors:** Mohsen Sadeghi

## Abstract

Peripheral membrane-associated proteins are known to accumulate on the surface of biomembranes as result of membrane-mediated interactions. For a pair of rotationally-symmetric curvature-inducing proteins, membrane mechanics at the low-temperature limit predicts pure repulsion. On the other hand, temperature-dependent entropic forces arise between pairs of stiff-binding proteins suppressing membrane fluctuations. These Casimir-like interactions have thus been suggested as candidates for attractive force leading to aggregation. With dense assemblies of peripheral proteins on the membrane, both these abstractions encounter multi-body complications. Here, we make use of a particle-based membrane model augmented with flexible peripheral proteins to quantify purely membrane-mediated interactions and investigate their underlying nature. We introduce a continuous reaction coordinate corresponding to the progression of protein aggregation. We obtain free energy and entropy landscapes for different surface concentrations along this reaction coordinate. In parallel, we investigate time-dependent estimates of membrane entropy corresponding to membrane undulations and coarse-grained tilt field and how they also change dynamically with protein aggregation. Congruent outcomes of the two approaches point to the conclusion that for low surface concentrations, interactions with an entropic nature may drive the aggregation. But at high concentrations, energetic contributions due to concerted membrane deformation by protein clusters are dominant.

---

Biomembranes are physical and chemical boundaries of cells and many of their organelles, as well as quasi-two-dimensional universes of encounters and actions behind a vast variety of vital functions [1]. Many aspects of membrane dynamics, configuration, topology, and even composition are actively controlled by membrane-associated proteins, facilitating, among others, cellular motility [2], proliferation [3], response to stimuli [4], trafficking and endocytosis [5–8]. This renders the understanding of localization, organization and action of these proteins of utmost importance. Peripheral proteins are a prominent class of membrane proteins, which can, through curvature-generation, contribute to membrane remodeling [9, 10]. The actual mechanism with which these proteins can sense or induce membrane curvature may vary [11–14], but the important commonality is how the concerted action of a collection of these proteins can lead to macroscopic membrane deformations [15, 16]. As an example, it was established early on that the internalization of Cholera toxin from cell membrane into endoplasmic reticulum can progress with or without relying on clathrin- or caveolae-mediated endocytosis [17]. Or when bound to glycolipids on the outer leaflet of giant unilamellar vesicles (GUV’s), the pentagonal subunit B of Shiga toxin (STxB) was shown to induce invaginations, and long tubules, in the absence of active cellular machinery [18]. This rather elegant entry mechanism, which has promoted STxB as a model for glycolipid-dependent endocytosis [19], has also inspired using it as a vehicle for drug delivery and anti-cancer therapy purposes [20–22]. The fact that membrane remodeling leading to internalization of these toxins can progress without relying on energy-consuming cell machinery has fueled the research into understanding the underlying protein-membrane mechanical interplay [15, 23, 24].

Organization of peripheral proteins is a complex and multi-scale phenomenon. Focusing on peripheral proteins inducing isotropic local curvature, effects ranging from lipid compression [14] and lipid phase separation [25] to active cytoskeletal forces [26] are considered to play a role. Focusing on length-scales at which the membrane can be modeled as a continuous surface, the free energy density corresponding to membrane deformations very well conforms with the Helfrich function 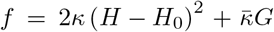, where *H, G* and *H*_0_ are the mean, Gaussian and spontaneous curvatures, and *κ* and 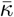 are the attributed elastic constants [27–31]. Utilizing this model, and minimizing total free energy in the presence of two inclusions with isotropic curvature of the same sign predicts “curvature-mediated” repulsion [32–34]. Similar repulsion has also been shown to sort partially budded membrane domains [35]. On the other hand, a dense assembly of curvature-inducing proteins can minimize energy with collective membrane deformations, and even result in apparent attraction [36]. Attractive interaction is also possible when inclusions possess anisotropic curvatures [32, 37–39]. On the other hand, temperature-dependent “fluctuation-mediated” interactions have been implicated as the source of long-range attraction [23, 34, 40–44]. These thermal Casimir-like forces push proteins that suppress membrane fluctuations together to minimize their adverse entropic effect. Simulations focusing on these interactions have confirmed their effectiveness in protein organization [24, 45].

Theoretical models and simulations in simplified scenarios that focus on certain type of membrane-mediated interactions are illuminating, but lack the generality in a more realistic setup where a dynamic membrane crowded with large numbers of peripheral proteins is evolving at timescales appropriate to the corresponding slow dynamics. The multi-scale nature of the problem connotes the usual trade-off between size and details, making it challenging to propose a model that can capture several aspects of peripheral protein interactions and organization. This spatiotemporal range of interest is very well covered by the so-called interacting particle reaction-diffusion (iPRD) models [46–49]. Proper to these models, we have recently developed an ultra coarse-grained membrane model [50, 51], with an accompanying hydrodynamic coupling method [52], which offers a remarkable balance between mimicking membrane mechanics and dynamics at the large scale and offering the possibility of small-scale effects, including flexible peripheral proteins inducing local curvatures [53, 54]. We implemented these membrane-associated proteins as particles tagged in the model to carry a masked force field as they freely diffused along the membrane. We took painstaking care to prevent the peripheral proteins to experience direct interactions as result of force field masking (see [54] and its Supplementary Information). The proteins aggregated on the membrane nonetheless, and the stiffest among them could form macroscopic clusters, given enough surface concentration, leading us to the conclusion that forces mediated by the membrane were responsible for this organization. Here, via similar large-scale simulations, we look into the actual origin of these forces. Specifically, we investigate the duality between energetic curvature-mediated and entropic fluctuation-mediated interactions in details. Using a liquid-mixture model, we first quantify pairwise membrane-mediated interactions and demonstrate the effect of protein stiffness and concentration. We subsequently use the continuous reaction coordinate highlighting protein aggregation to investigate the entropy landscape as a function of protein aggregation at different surface concentrations. We compare these entropies with two-dimensional configurational entropy due to surface distribution of proteins to show where aggregation enhances or diminishes overall entropy. Finally, we propose a method for direct estimation of entropy corresponding to fluctuations in membrane amplitude and lipid tilt field from simulation trajectories, and successfully repeat the same analysis thereon.

## RESULTS

### Magnitude of membrane-mediated interactions

We have simulated membrane patches containing peripheral proteins of intrinsic curvature of 0.08 nm^*−*1^, with the stiffness of 50, 100, and 200 MPa, and surface concentrations in the range 2 – 8 nmol m^*−*2^ using our mesoscopic modeling framework (details in Methods and [54]). The model allows for observation and measurement of the emergent membrane-mediated interactions between these proteins. We employ the statistical mechanics of liquid mixtures to estimate the effective interactions between pairs of the constituents. We consider a two-dimensional fluid, consisting of either pure membrane (m) or protein-bound (p) particles, and respectively denote the effective pairwise interactions as *u*_mm_, *u*_mp_, and *u*_pp_. We estimate these effective interactions from the set of all pair-correlation functions (details in Methods section “Statistical mechanics of liquid mixtures”). The interaction potential *u*_pp_ also contains contributions from the underlying membrane particles. To correct for this, we have simply calculated the net membrane mediated interaction as *u*_pp_ − *u*_mm_ (Fig. 1B).

**Figure 1:**
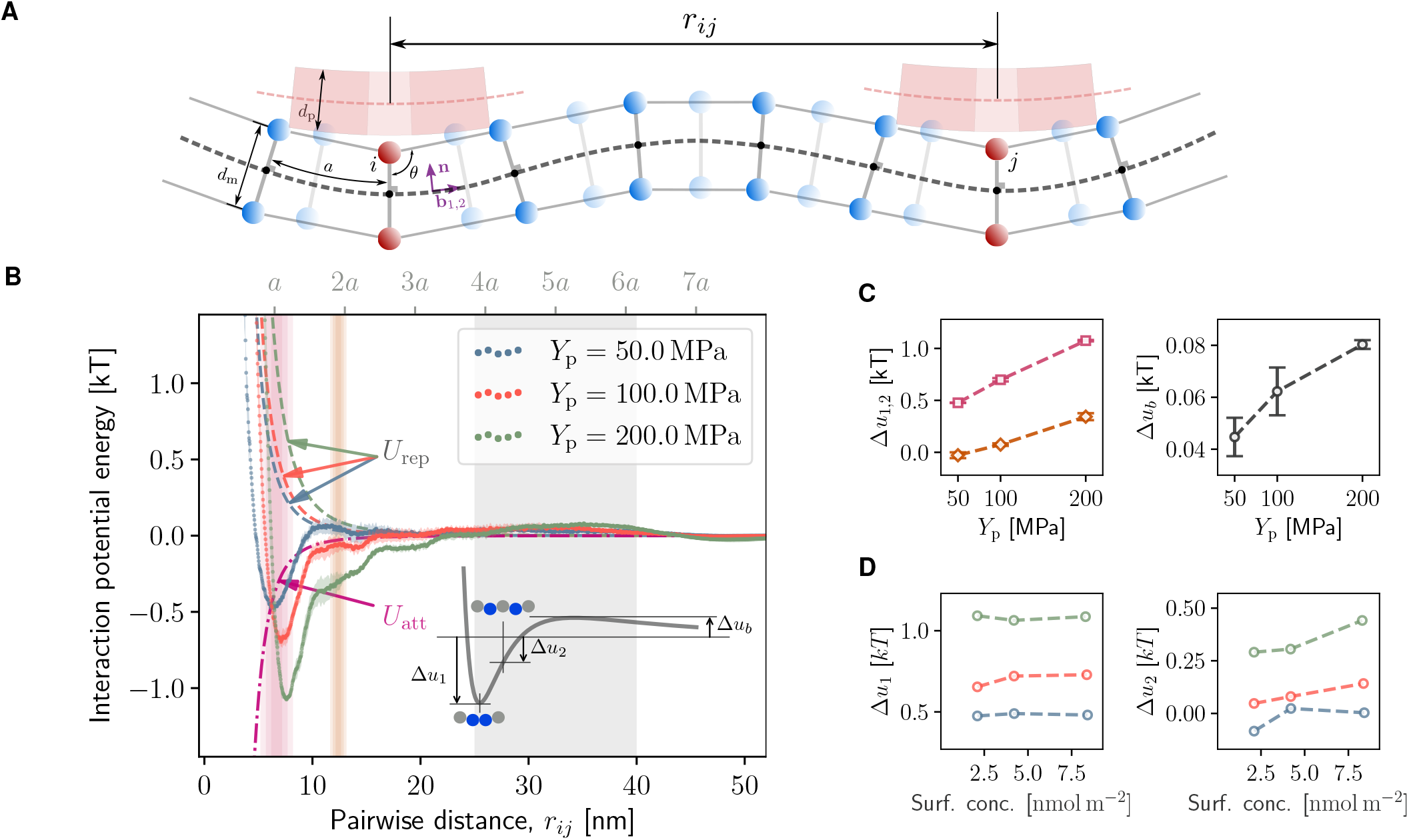
Membrane-mediated interactions in the dynamic model of membrane and peripheral proteins: **(A)** Mesoscopic membrane model with selected particles “tagged” as supporting peripheral membrane-bound proteins. Where a protein is bound to the membrane, a locally modified force field is applied (masking of the force field). Force field parameters are optimized to reproduce the Helfrich energy density for regular particles, plus an additional strain energy corresponding to the deformation of peripheral proteins for tagged particles (Methods section “Particle-based model of membrane and peripheral proteins”). **(B)** Interaction potential as a function of inter-particle distance, for peripheral proteins with the given stiffness, inferred from simulations using the liquid-mixture model (Methods section “Statistical mechanics of liquid mixtures”). Repulsive curvature-mediated interaction *U*_rep_, as well as the fluctuation-mediated attractive interaction *U*_att_ are plotted for comparison (Eqs. (1) and (2)). Shaded pink and orange regions correspond to first and second neighborhood of particles, determined from position of peaks in the pair-correlation functions. The width of both regions is characterized by the shown six standard deviations from the mean distance. Gray shaded region designate where the heights of the repulsive barriers have been sampled. **(C)** Effect of the protein stiffness on the shape and strength of the interaction potential. Well-depth, Δ*u*_1,2_, and barrier height, Δ*u*_b_, are measured according to the inset schematic shown in (B). **(D)** Effect of the protein surface concentration on the attraction between a protein and its first and secondary neighboring proteins.

We consider two prominent continuum-based models for comparison: (i) repulsive mean-field interactions between symmetrically curved regions, either from peripheral proteins or from stiff inclusions in the membrane [32–34]. These interactions stem from minimizing membrane elastic energy, neglecting fluctuations, thus corresponding to equilibrium at zero temperature [40]. For a pair of particles with the separation *r*, this interaction is approximated in the leading power by [34, 40, 44],

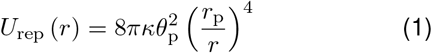

where *θ*_p_ and *r*_p_ are the contact angle and the radius of the curved region and *κ* is the bending rigidity of the membrane. (ii) attractive entropic forces emerging when the binding is stiff enough to locally suppress membrane fluctuations at non-vanishing temperature (the so-called thermal Casimir effect). An approximate expression for the potential of this attractive interaction is [34, 40, 44],

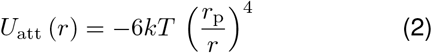

Membrane-mediated interaction potentials that we obtained from the simulations show significant attraction at close range (Figs. 1B). For the stiffest proteins considered, this attraction is present even up to particle separations of 3*a* where *a* is the lattice parameter of the model (Figs. 1A and 1B). Attraction strength (depth of the potential well) for the first and second neighbors (Δ*u*_1_ and Δ*u*_2_, respectively) is proportional to the protein stiffness (Figs. 1B and 1C). Yet, looking at how the surface concentration of proteins affects the attraction (Fig. 1D) reveals non-trivial dependence, especially for the second-neighbor interactions. This points to the shortcoming of pairwise interaction model underlying the liquid-mixture model and reveals weak multi-body contributions.

Higher protein stiffness results in more pronounced curvatures [54], and subsequently closer possible approach of tagged particles in two dimensions (Fig. 1A). Yet, with the increase in protein stiffness, the close-range repulsion occurs at larger distances, evidently beyond the unperturbed lattice parameter, *a* (Fig. 1B). We thus conclude that aside from the steric repulsion of particles, which is similar for all cases, a curvature-mediated repulsion is also present.

We also observe a small repulsive barrier at the distance of ∼ 5*a*, which is not predicted by either of the two models (Fig. 1B). There is a clear relation between the barrier height, Δ*u*_*b*_, and protein stiffness (Fig. 1C) which implies a long-range curvature-mediated effect.

### Entropy of protein aggregation from histogram analysis

We have performed equilibrium simulations with different surface concentration of peripheral proteins with the stiffness of 200 MPa and have used a reaction co-ordinate *q* to be a continuous descriptor of aggregation. This reaction coordinate is defined as the mean of inverse pairwise distances between bound proteins, with larger *q* values implying stronger aggregation (Figs. 2A and 2B, Methods section “Entropy Estimation” and [54]). We have affirmed the sensitivity of *q* to protein aggregation by obtaining clear separation of distributions for states that differ minutely in mean cluster size (Fig. 2C).

**Figure 2:**
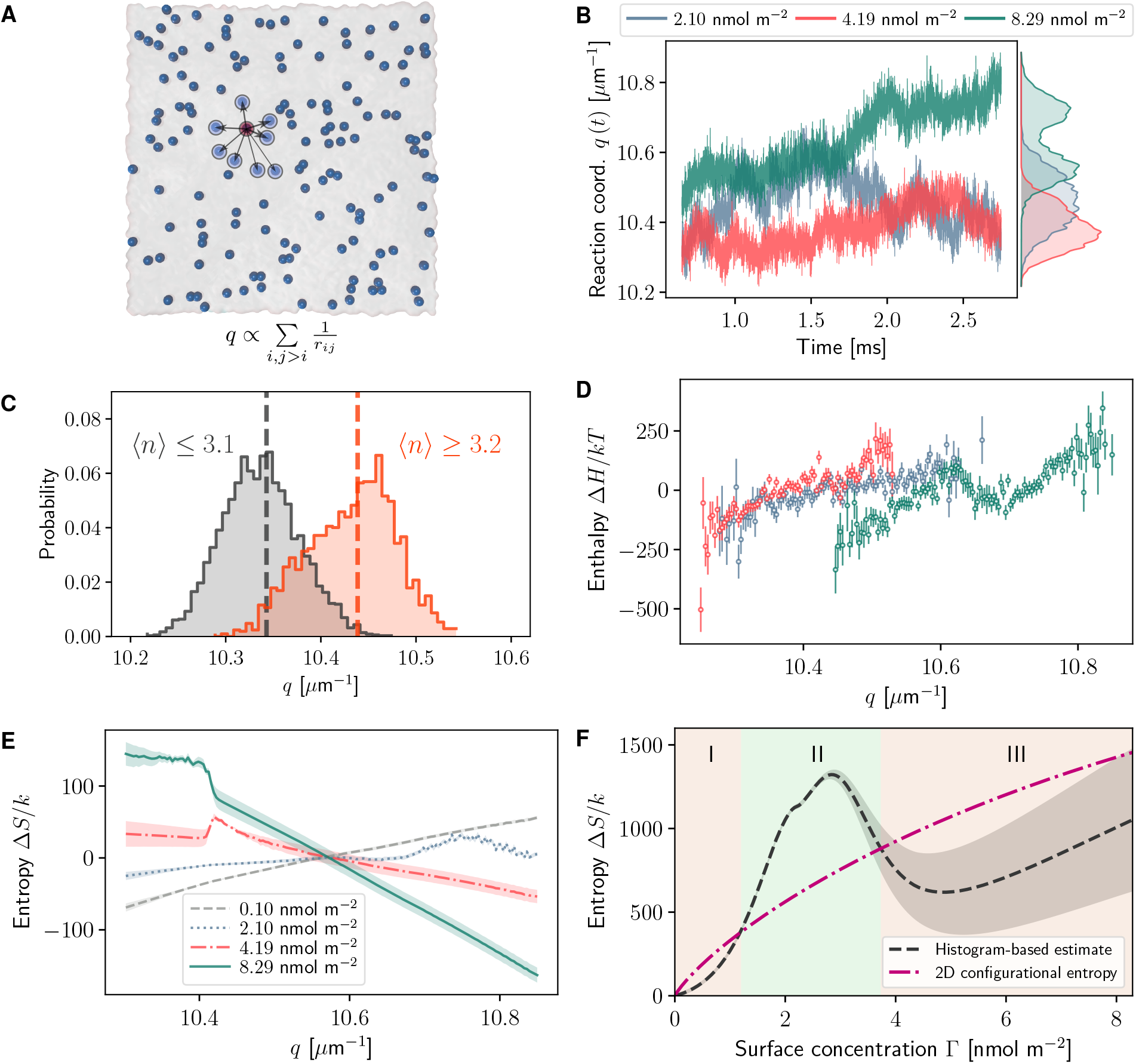
Entropy change due to protein aggregation studied via histogram-based estimates based on a continuous reaction coordinate: **(A)** Definition of the reaction coordinate *q* corresponding to the degree of aggregation, based on the pairwise distances between peripheral proteins. **(B)** Time evolution of the reaction coordinate for simulations with the given surface concentration of proteins. **(C)** Histograms of reaction coordinate, split for frames in which the mean cluster size, ⟨*n*⟩, complies with the given criteria. **(D)** Enthalpy, binned along the reaction coordinate *q*. Colors correspond to different surface concentrations, following the legend of panel (B). **(E)** Entropy change as a function of protein aggregation, which is indicated by the reaction coordinate, *q*, for the given protein surface concentrations. **(F)** Mean entropy as a global function of surface concentration of proteins. The two-dimensional configurational entropy of proteins (Eq. (10)) is also shown for comparison.

We have obtained histograms of states along *q* and have used reweighting based on a model of biased interactions to come up with continuous free energy and entropy landscapes (Methods section “Entropy Estimation”). We observe the entropy as a function of *q* to have a concentration-dependent behavior (Fig. 2E). For low surface concentrations of 0.10 nmol m^*−*2^ and 2.10 nmol m^*−*2^, the entropy increases with *q*, compliant with entropic forces pushing toward more aggregation. But as concentration increases to 4.19 nmol m^*−*2^ and 8.29 nmol m^*−*2^, this trend is reversed, with more aggregation lowering the entropy (Fig. 2E). The histograms of enthalpy along *q* reveals the energy to increase with *q* for low concentrations (Fig. 2D), but at the high concentrations, local energy minima explaining the observed aggregated states are present (Figs. 2D and 2B).

Calculating the expectation value of the estimated entropy along *q* yields its mean value as a continuous function of surface concentration, Γ (Fig. 2F). We found it greatly beneficial to include a measure of two-dimensional configurational entropy of indistinguishable particles with the same surface concentration (Eq. (10) and Fig. 2F). Firstly because the numerical proximity of the two entropy estimates serves as a sanity check for the histogram-based estimation (Fig. 2F). Secondly, the “contest” between the two entropies introduces three distinct regimes (Fig. 2F): (I) histogram-based and configurational entropies are closely comparable at low Γ. Smaller histogram-based entropy might be due to insufficient sampling in this disperse regime. (II) At mid-range Γ, histogram-based entropy is remarkably higher than the 2D configurational estimate. This is possible because the histogram-based estimate also includes membrane contributions. In this regime, the increased entropy suggests new microstates available through amenable membrane configurations. This also perfectly matches our observation of the dominance of entropic interactions for this range of surface concentrations. (III) At high Γ, histogram-based entropy falls significantly lower than the pure configurational estimate, highlighting the fact that stable clustered configurations in this regime prevent the exploration of combinatorial configuration possibilities. Lowered entropy must be compensated by favorable energetic interactions (Fig. 2D).

Using the differential form *dG* = −*SdT* +*Adγ* +*μ*_p_*dN*_p_, where *γ* and *A* are the membrane tension and surface area, and *N*_p_ and *μ*_p_ are copy number and chemical potential of the bound proteins, a Maxwell relation can be written as,

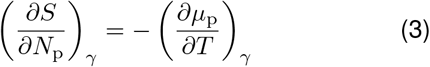

Thus, positive slope in the entropy-concentration plot of Fig. 2F means the chemical potential decreasing with temperature. Referring to the usual temperature dependence of entropic forces (e.g. Eq. (2)), this is equivalent to the increase in attractive entropic interactions.

### Direct estimation of membrane entropy

To gain better insight into the origin of the observed entropy variations, we directly estimate the entropy of the membrane from simulation trajectories, and observe its behavior as a function of protein aggregation. Generally, it is non-trivial to assign a quasi-equilibrium time-dependent estimate of entropy to a dynamical system [55–57]. Out main interest here is to observe the qualitative behavior of the entropy functional, and thus, we approach this simply via coarse-grained Gibbs entropy,

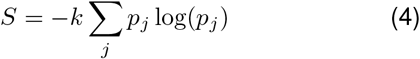

where the summation is carried out on all the states in an arbitrarily discretized configuration space and *p*_*j*_’s correspond to the probability of the system being found in the *j*-th discrete cell. We limit our measurement to two aspects of the entropy in this system: (i) Δ*S*_*h*_, the entropy assigned to membrane undulations, where the discretization is done in the Fourier space of fluctuation modes (Fig. 3A), and (ii) Δ*S*_**n**_, the orientational entropy of the tilt field, where the probability of states is measured based on *P*_2_ (cos *θ*), where *P*_2_ (*x*) is the Legendre polynomial of order 2, and *θ* is the local tilt angle (Figs. 1A and 3A). Counting the states, respectively based on the discretized fluctuation spectrum and the order parameter distribution, comprises a specific coarse-graining which renders the numerical values of the entropies thus obtained not comparable to other estimates. On the other hand, this specific accumulation of microstates results in sufficient sampling for each bin, even in a time-dependent evolving sense, when sampling is performed over sliding temporal windows (further details in Methods section “Entropy estimation”).

**Figure 3:**
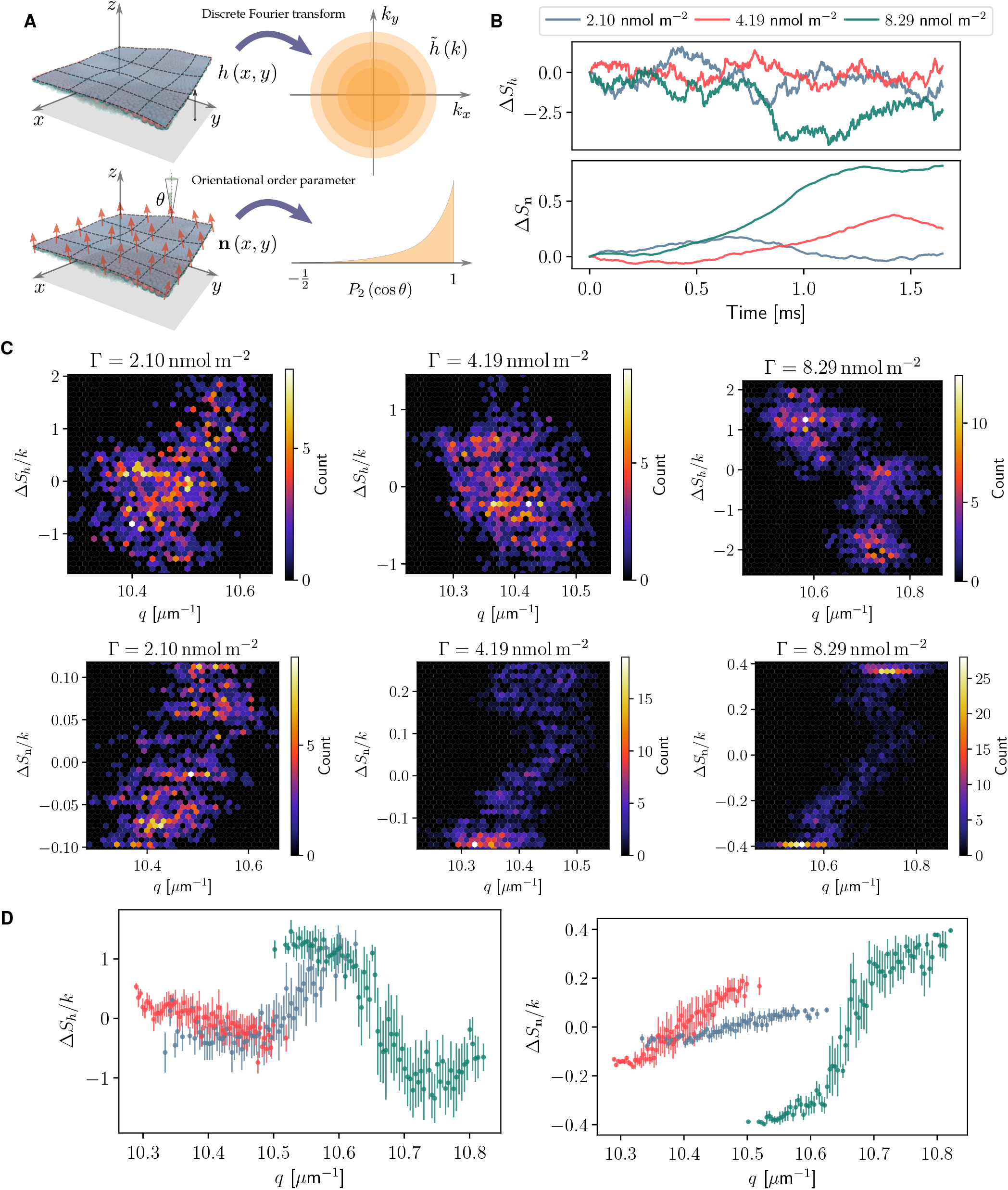
Entropy change due to protein aggregation based on direct estimation of membrane entropy from simulation trajectories. **(A)** Schematics depicting how the entropy of membrane undulations, Δ*S*_*h*_, (top) as well as the orientational entropy of the tilt field, Δ*S*_**n**_, (bottom) are estimated from dynamic histograms. **(B)** Time series of the two entropies for the given surface concentration of peripheral proteins. **(C)** Heat maps of estimated entropies when trajectory frames are also binned against the reaction coordinate *q*. **(D)** Undulation and tilt-field entropies as functions of the reaction coordinate *q* (colors correspond to the legend of panel (B)).

We observe that Δ*S*_*h*_ only increases with protein aggregation in the lowest concentration of Γ = 2.10 nmol m^*−*2^ (Figs. 3C and 3D). When higher concentrations are imposed (Γ = 4.19 and 8.29 nmol m^*−*2^), this entropy decreases with aggregation. The decrease becomes more drastic with higher concentrations (Figs. 3C and 3D). On the other hand, the tilt-field entropy, Δ*S*_**n**_, tends to increase with aggregation, independent of concentration (Figs. 3C and 3D). The slope of its increase is also directly proportional to surface concentration (Fig. 3D). Interestingly, with both Δ*S*_*h*_ and Δ*S*_**n**_, when Γ = 8.29 nmol m^*−*2^, we observe a clear separation of low and high entropy states (Fig. 3C).

## DISCUSSION

We have performed simulations using a particle-based membrane model with locally masked force field representing curvature-inducing bound proteins. This model predicts a complex aggregation behavior for these peripheral proteins based on their stiffness and surface concentration [54].

Using a liquid mixture model, we quantified the effective pairwise potential of membrane-mediated interactions between proteins. We observe a rather long-range attraction with a strength depending on protein stiffness. Surface concentration of particles has an effect on the pairwise interactions, especially when the pair of particles are not nearest neighbors (Fig. 1D). This suggests that multi-body effects are present. Idealized repulsion and attraction potentials originating from continuum models (Eqs. 1 and 2) are not applicable to the complex interaction potential observed. On the other hand, multi-body effects limit the applicability of the pairwise interaction model altogether. Application of more elaborate data-driven models to these interactions would thus be promising in gaining better insight as well as extrapolating the dynamics to larger timescales. We believe deep learning, especially via graph convolutional neural networks (GCNN) holds the key to such an investigation [58, 59].

We obtained entropy landscapes along a continuous aggregation reaction coordinate, and showed that entropy only increases with aggregation when the surface concentration is below a threshold. Comparison of mean entropy as a function of surface concentration revealed that only for concentrations below this threshold the estimated entropy can surpasses the 2D configurational entropy of proteins as indistinguishable particles on the membrane. We thus conclude that while entropic forces originated from membrane fluctuations exist and are apparently responsible for bringing proteins together in the low-concentration regime, they are overshadowed at higher concentrations. At high concentrations, there are enough random encounters between particles on the surface of the membrane that the energy minima due to collective membrane deformation become more influential.

To further corroborate these findings, we took an independent measurement via direct estimation of coarse-grained Gibbs entropy. We looked at the entropy of membrane undulation modes as well as orientational entropy of the tilt field. We found out that the generally larger undulation entropies point to the same trend as the histogram-based estimates. We also observed that the orientational entropy of the tilt field always favors aggregation. It is imaginable that in a model with a higher resolution representation of lipids, this entropy have a more pronounced effect. But in our model, and with the admittedly arbitrary partitioning of configuration space, the undulation entropies seem to be always dominant in magnitude, and resisting the aggregation at high concentrations.

We believe the dynamic membrane + protein model and the detailed analyses presented here can be greatly beneficial in investigating different scenarios for membrane-mediated collective action. Ever-advancing super-resolution microscopy [60] and optical spectroscopy methods [61] make it possible to look at organization of membrane-associated proteins in unprecedented details, while resolving membrane fluctuations at nanometer/microsecond range. This opens the door to direct quantitative comparison with experimental observation of membrane-mediated effects, and will pave the way for unraveling and controlling the underlying mechanisms.

## METHODS

### Particle-based model of membrane and peripheral proteins

The particle-based model of membrane and bound peripheral proteins used in this study is fully described in [54]. This model is an extension of our previously established dynamic membrane model [50–52]. We have used a lattice parameter of 6.5 nm for all the particles, comparable with the diameter of the STxB protein. We use a force field for describing nearest neighbor interactions in this model which allows a local masking where peripheral proteins are bound (Fig. 1A). We have utilized the combined energy density of membrane and the protein in conjunction with parameter-space optimization to obtain the global force field parameters [54] (Fig. 1A).

### Statistical mechanics of liquid mixtures

We assume rotational isotropy in the 2D fluid, and consider the pair-correlation function between the species *α* and *β* as *g*_*αβ*_ (*r*). Defining *h*_*αβ*_ (*r*) = *g*_*αβ*_ (*r*) − 1, the so-called direct correlation function, *c*_*αβ*_ (*r*), is defined implicitly [62],

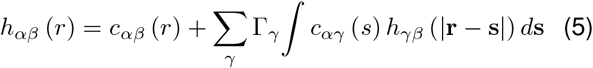

where Γ’s denote surface densities. Eq. (5) translates to

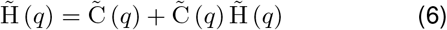

where 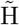 and 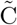 are 2 × 2 matrices with elements 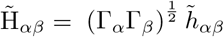 and 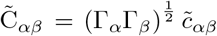 and 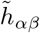 and 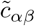 are two-dimensional Fourier transforms of *h*_*αβ*_ and *c*_*α,β*_. Eq. (6) is formally known as the *Ornstein-Zernike* relation [62, 63]. Several “closure” relations are available which tie *h*_*αβ*_, *c*_*αβ*_, and *g*_*αβ*_ via a pairwise potential *u*_*αβ*_ between *α* and *β*. We have employed the convoluted hypernetted-chain (CHNC) relation [64],

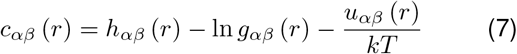

The procedure for obtaining the potentials *u*_mm_, *u*_mp_, and most importantly, *u*_pp_, is as follows: we calculate discretized estimates of the three pair correlation functions, *g*_mm_ (*r*), *g*_mp_ (*r*), and *g*_pp_ (*r*), from the simulation trajectories, and substitute numerically obtained 2D Fourier transforms into Eq. (6) to obtain *c*_*αβ*_’s and subsequently *u*_*αβ*_’s from Eq. (7).

### Entropy estimation

As is implied in Fig. 2A, the reaction coordinate *q* is defined based on the inverse pairwise distances between proteins [54],

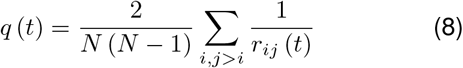

Each simulation with a fixed surface concentration yields a histogram of microstates along *q*. Because of the fact the implementation of the masked force field only modifies the interaction potential without changing the number of particles, we make the assumption that the Hamiltonian is biased as ℋ= ℋ_0_ + *ξ f* (*q*), with *ξ* proportional to the surface concentration. Based on this assumption, the multiple histogram method of Ferrenberg and Swendsen [65] (which is the precursor to the well-known Weighted Histogram Analysis Method (WHAM) [65, 66]) can readily be applied to combine the histograms from different simulations.

We assume that each simulation, in which a different number of particles have been tagged as proteins, is biased with a Hamiltonian of the form ℋ= ℋ_0_ + *ξ f* (*q*). A justification for this assumption is given in [54]. We have chosen polynomials of degree 4 as the function *f* (*q*), and have fitted to randomly sub-sampled energy data to obtain an error estimate. Application of multiple histogram reweighting yields the free energy landscape along *q* for different values of *ξ*, which correlate with surface density of proteins. Once the free energy change, Δ*G*, is available, we can estimate the entropy chage as,

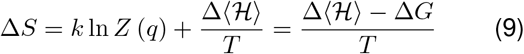

where *Z* (*q*) is the isothermal-isotension partition function.

For estimating the two dimensional configurational entropy of proteins spattered on the membrane surface, we have simply assumed that they can occupy any of the available sites on the particle-based membrane (Fig. 1A). A simple combinatorial assessment followed by the Stirling’s formula yields,

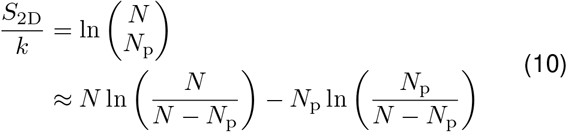

where *N* and *N*_p_ are the total number of particles in the upper leaflet and the number of proteins, respectively.

We have obtained the undulation entropy, Δ*S*_*h*_, by first mapping the height of membrane particles to a regular 2D grid using bilinear interpolation, i.e. to get a regular sampling of the height function, *h* (*x, y*). Following a spatial fast Fourier transform for each time frame, we have discretized 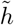(k) in the frequency domain based on the length of the wave-vector, *k* (Fig. 3A) and obtained a sliding-window estimate of the probability distribution from the histogram of states. For the tilt-field entropy, Δ*S*_**n**_, the discretization is achieved in real space based on samples of the orientational order parameter *P*_2_ (cos *θ*) (Fig. 3A). For both entropies, point values has been estimated on sliding-window samples over a 0.5 ms temporal window.

## SOFTWARE AVAILABILITY

All the in-house developed software used in this study are available from the author upon reasonable request.

## DATA AVAILABILITY

The data that support the findings of this study are available from the author upon reasonable request.

## ACKNOWLEDGEMENTS

The author is especially grateful to Frank Noé (FU Berlin) for constant support and valuable comments and discussions. We also acknowledge helpful comments from Thomas Weikl (MPI for colloids and interfaces) and Patricia Bassereau (Institut Curie). This research has been funded by Deutsche Forschungsgemeinschaft (DFG) through grants SFB 958/A04 and SFB 1114/C03.

